# Facility ownership and mortality among older adults residing in care homes

**DOI:** 10.1101/319459

**Authors:** Javier Damián, Roberto Pastor-Barriuso, Fernando José García-López, Ana Ruigómez, Pablo Martínez-Martín, Jesús de Pedro-Cuesta

## Abstract

**Background and Objectives:** Nursing or care home characteristics may have a long-term impact on the residents’ mortality risks that has not been studied previously. The study’s main objective was to assess the association between facility ownership and long-term, all-cause mortality. *Research Design and Methods*. We conducted a mortality follow-up study on a cohort of 611 nursing-home residents in the city Madrid, Spain, from their 1998–1999 baseline interviews up to September 2013. Residents lived in three types of facilities: public, subsidized and private, which were also sub-classified according to size (number of beds). Residents’ information was collected by interviewing the residents themselves, their caregivers and facility physicians. We used time-to-event multivariable models and inverse probability weighting to estimate standardized mortality risk differences.

**Results:** After a 3728 person-year follow-up (median/maximum of 4.8/15.2 years), 519 participants had died. In fully-adjusted models, the standardized mortality risk difference at 5 years of follow-up between large-sized public facilities and medium-sized private facilities was 18.9% (95% confidence interval [CI]: −33.4 to −4.5%), with a median survival (95% CI) of 3.6 (0.5 to 6.8) additional years. The fully-standardized 5-year mortality difference (95% CIs) comparing for-profit private facilities with not-for-profit public institutions was −15.1% (−31.1% to 0.9%), and the fully-standardized median survival difference (95% CIs) was 3.0 (-1.7 to 7.7) years.

**Discussion and Implications:** These results are highly compatible with an effect on the long-term mortality risk for residents due to factors associated with the ownership of their facilities.

## Introduction

Long-term care for older adults is evolving rapidly into many diverse alternatives but nursing homes still are an essential component of the current sector [1] and the need for these facilities is expected to grow along with countries’ aging populations. In a few years the high life-expectancy in many countries will lead to frequent instances of older adults caring for their parents – older people caring for even older people. In this situation nursing home alternatives are expected to increase in number and variability.

Mortality is an indicator of quality of care in nursing homes, but with complex determinants [1–3]. Studies focusing on mortality in care homes associated with ownership come mostly from North America, mainly from the USA [4–13] (see Table S1 at supplementary material). In addition, most investigations are focused on short-term mortality. Some of these [9, 13] began a short follow-up of residents just after admission; consequently, the influence of facility characteristics on mortality will be limited, and health and mortality can be excessively affected by pre-admission factors. Most research has focused on ownership using the contrasting for-profit/not-for-profit criteria. In country Spain, however, the classification private, public and subsidized is common. In this study, the baseline sampling used a stratification of the population based on whether the resident was publicly supported or not, in particular whether s/he lived in a public or subsidized facility, or in a private one; consequently, we opted for using a private/public classification, although we have also examined the for-profit/not-for-profit schema (see supplementary material for definitions used in the present work). Hence, the main objective of this study was to measure the long-term mortality risk of residents according to the type of facility where they live, with particular regard to ownership.

## Methods

### Study population

This cohort study used mortality follow-up data from a baseline survey conducted from June 1998 through June 1999 in a representative sample of residents aged 65 years or older in residential and nursing homes in Madrid, Spain. Study participants were selected through stratified cluster sampling, including one stratum with 47 public or subsidized (privately owned but publicly funded) nursing homes and another stratum with 139 private institutions. We initially selected 25 public/subsidized and 30 private institutions with probability proportional to their size (range 16 to 620 beds), and then sampled 10 men and 10 women from each selected public/subsidized facility and five men and five women from each selected private facility by means of systematic sampling with random start, using their complete alphabetical list of residents. Four private institutions declined to participate (totaling 40 sample participants) and 45 additional residents could not be selected due to prolonged absence or refusal, leading to an overall response rate of 89% (715 out of 800 sample residents). Of the 45 missing, thirty nine participants could be randomly substituted with residents of the same facility and sex, yielding a total of 754 residents. As a result of this design, residents in public/subsidized facilities and men were oversampled and hence sampling weights were assigned to study participants as the inverse of their selection probabilities.

The Carlos III Institute of Health Institutional Review Board approved the study. Informed consent was obtained verbally from all study participants or their next of kin and documented. This investigation was conducted according to the principles expressed in the Declaration of Helsinki.

### Baseline data collection

Structured questionnaires were administered verbally by trained geriatricians or residents in geriatrics, to all selected residents, their main caregivers, and the facility physicians or nurses – in order to collect baseline data on sociodemographic characteristics, medical conditions, and functional dependency. Age, sex, educational level (less than primary [8 years], or primary or more), and length of stay in the nursing home were obtained by interviewing residents or a proxy. Chronic medical conditions – including cancer, chronic obstructive pulmonary disease, arrhythmias, ischemic heart disease, congestive heart failure, peripheral arterial disease, stroke, hypertension, diabetes, anemia, Alzheimer’s disease, other dementias, Parkinson’s disease, epilepsy, depression, anxiety disorders, and arthritis – were ascertained by interviewing facility physicians (or nurses for 8% of residents) who had access to medical histories. The number of chronic conditions other than dementia was computed and categorized into 0–2 and ≥3 diseases. Functional dependency – ability to perform basic activities of daily living – was assessed by interviewing the residents (59%) or their main caregivers (41%) using the modified Barthel index [14]. Based on previously proposed cut-offs [14], residents were classified as being functionally independent or mildly dependent (91–100 points), moderately dependent (61–90 points), and severely or totally dependent (0–60 points).

### Mortality ascertainment during follow-up

Study participants were followed up until 15 September 2013. Mortality was ascertained by requesting updated data on residents’ vital status from the participating facilities and through computerized linkage to the Spanish National Death Index, which includes all deaths registered in country Spain since 1987 [15]. Residents contributed follow-up time from their 1998–1999 baseline interview until death, age 105 years, or 15 September 2013, whichever came first.

### Statistical analysis

In primary analyses facilities were classified according to their ownership (public/subsidized, or private) and size (<100, 100–299, or ≥300 beds) as large-sized public, medium-sized public/subsidized, medium-sized private, and small-sized private institutions. Preliminary analyses showed similar mortality among residents in medium-sized public and subsidized facilities, so that both types of facilities were aggregated into a single category. For comparison with previous studies, facilities were categorized in secondary analyses according to ownership and profit status into: not-for-profit public, for-profit subsidized, not-for-profit private, and for-profit private institutions.

The cumulative all-cause mortality function over time for each type of facility was standardized to the weighted distribution of selected confounders in the overall study population of institutionalized residents by using inverse probability weighting [16–18]. We estimated each resident’s population conditional probability of being in its own type of facility given the observed confounders (see below) by fitting sampling-weighted multinomial logistic regression models with type of facility as outcome and the selected confounders as explanatory variables. Standardization weights were then calculated as the inverse of these estimated conditional probabilities of facility type, further stabilized by multiplying them by each of the four types of facility’s marginal proportions (sampling weighted) [16]. Finally, combined weights were assigned to study participants as the product of their sampling weights – which corrected the sample for selection bias to represent the population – and the stabilized weights – which adjusted for confounding [17].

Two sets of models were used. The first included baseline sociodemographic characteristics, such as age (65–74, 75–79, 80–84, 85–89, or ≥90 years), sex (woman or man), educational level (less than primary, or primary or more), and length of stay in the nursing home (<3 or ≥3 years); the second set of models further added dementia (yes or no), number of chronic conditions other than dementia (0–2 or ≥3), and functional dependency (no/mild, moderate, or severe/total). The mean combined weights were 1.00 (range, 0.16–4.45) for sociodemographic, and 0.99 (range, 0.15–7.86) for complete models. This weighting procedure achieved an effective standardization, since the fully-weighted distributions of baseline confounders were nearly identical across types of facility and closely matched their sampling-weighted distributions in the overall institutionalized population (data not shown).

### Mortality analysis (more detailed description in supplementary material)

For mortality risk analyses, we used Kaplan-Meier methods and spline-based survival models [19] weighted by the combined weights and stratified by type of facility. We used these models to estimate standardized differences in cumulative mortality at 2, 5, and 10 years of follow-up for each type of facility compared with large-sized public facilities (as a reference category), with 95% confidence intervals (CIs). In addition to risk differences, we estimated standardized differences in median follow-up times (50% cumulative mortality) and their 95% CIs. We evaluated homogeneity in risk differences across pre-specified subgroups of residents defined by baseline age (65–84 or ≥85 years), sex (woman or man), educational level (less than primary or primary or more), length of stay (<3 or ≥3 years), dementia (yes or no), number of chronic conditions other than dementia (0−2 or ≥3), and functional dependency (no/mild or moderate/severe/total) by fitting spline-based survival models weighted by combined weights and stratified by type of facility and resident subgroup.

Statistical analyses were performed using the *stpm* command in Stata, version 14 [20] and graphics were produced in R, version 3 [21].

## Results

Of the 754 participants in the baseline survey, we excluded 88 residents (12%) with missing information on any baseline covariate and 55 residents (7%) with unknown vital status at the end of follow-up, thus leaving a final cohort of 611 residents. Residents in private facilities had higher educational levels than those in public/subsidized facilities. Those in large-sized public facilities had longer stays and more chronic conditions, but lower degrees of functional dependency at baseline than those in any other type of facility (Table 1).

**Table 1.** Baseline characteristics of institutionalized residents by type of facility in city Madrid, Spain, 1998–1999.^a^

| Characteristic | Overall | Facility |  |  |  | *P*<br>value<sup>b</sup> |
| --- | --- | --- | --- | --- | --- | --- |
|  |  | Large-sized<br>public | Medium-sized<br>public/subsi<br>dized | Medium-sized<br>private | Small-sized<br>private |  |
| No. of residents | 611 (100) | 276 (38.3) | 156 (20.4) | 78 (18.3) | 101 (23.0) |  |
| Age (years) | 83.4 (7.3) | 84.1 (7.4) | 84.1 (6.9) | 83.1 (7.4) | 82.1 (7.7) | 0.12 |
| Sex |  |  |  |  |  | 0.15 |
| Women | 328 (74.5) | 146 (73.9) | 72 (68.4) | 50 (79.9) | 60 (76.6) |  |
| Men | 283 (25.5) | 130 (26.1) | 84 (31.6) | 28 (20.1) | 41 (23.4) |  |
| Educational level |  |  |  |  |  | <0.001 |
| Less than<br>primary | 280 (42.3) | 147 (54.2) | 94 (60.6) | 16 (21.2) | 23 (23.0) |  |
| Primary or more | 331 (57.7) | 129 (45.8) | 62 (39.4) | 62 (78.8) | 78 (77.0) |  |
| Length of stay<br>(years) |  |  |  |  |  | <0.001 |
| <3 | 287 (47.0) | 78 (25.7) | 106 (69.7) | 43 (52.9) | 60 (57.6) |  |
| ≥3 | 324 (53.0) | 198 (74.3) | 50 (30.3) | 35 (47.1) | 41 (42.4) |  |
| Dementia |  |  |  |  |  | 0.97 |
| No | 468 (75.1) | 213 (74.5) | 123 (76.7) | 59 (75.8) | 73 (74.1) |  |
| Yes | 143 (24.9) | 63 (25.5) | 33 (23.3) | 19 (24.2) | 28 (25.9) |  |
| No. of chronic<br>conditions<sup>c</sup> |  |  |  |  |  | 0.002 |
| 0–2 | 296 (49.2) | 114 (38.5) | 83 (53.0) | 41 (53.2) | 58 (60.6) |  |
| ≥3 | 315 (50.8) | 162 (61.5) | 73 (47.0) | 37 (46.8) | 43 (39.4) |  |
| Functional dependency |  |  |  |  |  | 0.003 |
| No/mild | 338 (51.4) | 184 (63.2) | 78 (47.7) | 37 (45.1) | 39 (39.9) |  |
| Moderate | 137 (24.2) | 54 (21.2) | 41 (26.3) | 16 (22.9) | 26 (28.2) |  |
| Severe/total | 136 (24.4) | 38 (15.6) | 37 (26.0) | 25 (32.0) | 36 (31.9) |  |

<sup>a</sup> Unweighted counts (sampling-weighted percentages).
<sup>b</sup> *P* value for homogeneity of sampling-weighted proportions across types of facility.
<sup>c</sup> Number of chronic conditions other than Alzheimer's disease and other dementias.

A total of 519 participants died during the 3,728 person-years of follow-up (median [interquartile range] follow-up of 4.8 [2.0–9.5] years), with an overall mortality rate of 133 per 1000 person-years. After standardization to the overall weighted population distribution of age, sex, educational level, length of stay, dementia, number of chronic conditions, and functional dependency, all-cause mortality was substantially lower among residents in medium-sized private facilities than in other types of institutions (Fig 1). Compared with residents in large-sized public facilities, the standardized mortality risk differences at 5 and 10 years of follow-up were −6.1% and −1.5% for residents in medium-sized public/subsidized facilities, −18.9% and – 17.7% for those in medium-sized private facilities, and −12.9% and −5.3% for those in small-sized private facilities (Table 2). Similarly, the standardized differences in the median survival time comparing residents with those in large-sized public institutions were 0.9 years in medium-sized public/subsidized, 3.6 in medium-sized private, and 1.9 in small-sized private facilities respectively (Table 3).

**Fig 1.**
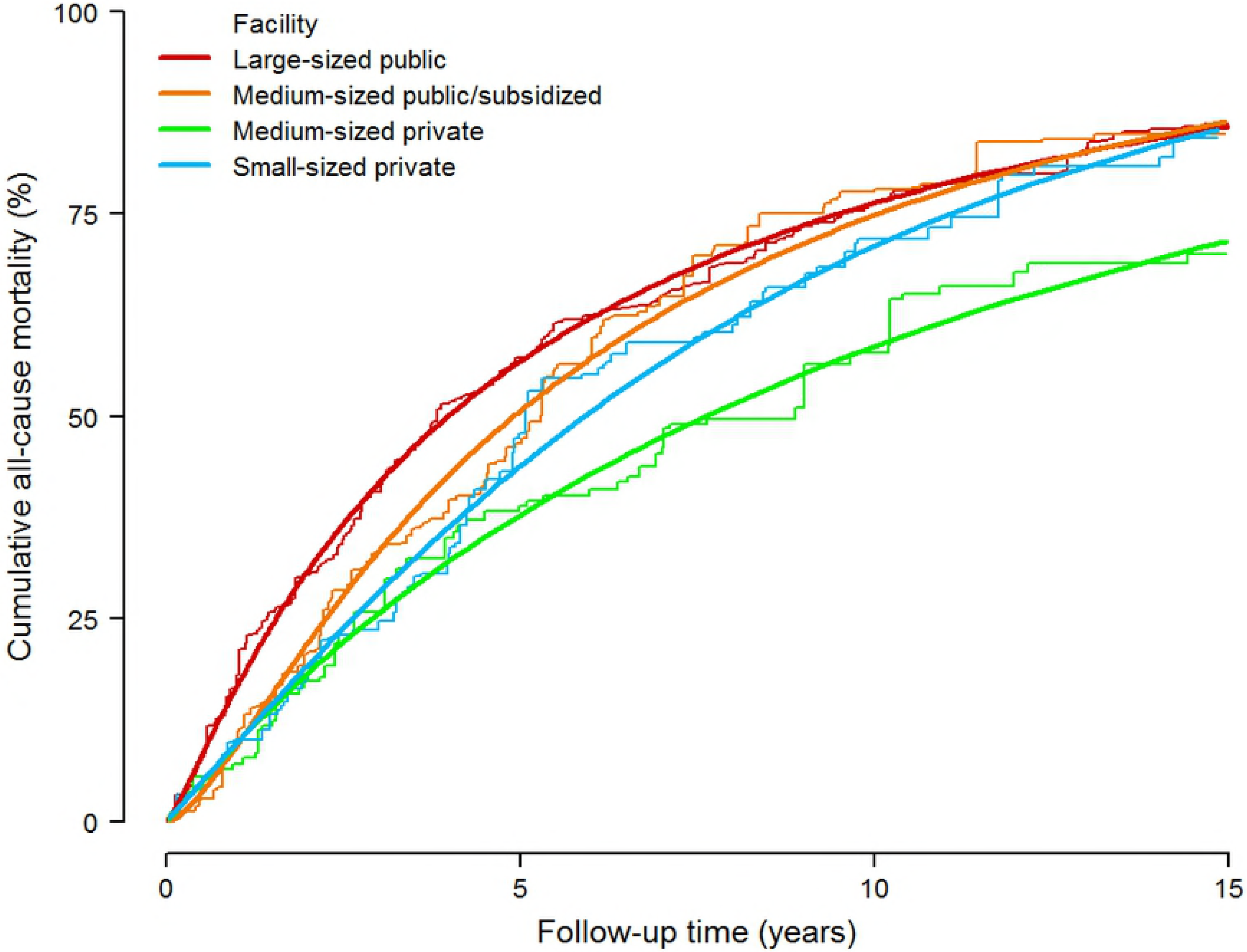
Standardized cumulative all-cause mortality by type of facility among institutionalized residents in Madrid, Spain. Parametric cumulative mortality curves (smooth lines) were estimated from a spline-based survival model and nonparametric cumulative mortality curves (step functions) from Kaplan-Meier methods, both weighted by combined inverse probability weights and stratified by type of facility. Combined weights were used to standardize cumulative mortality curves in each type of facility to the weighted distribution of baseline confounders in the overall institutionalized population, including age (65–74, 75–79, 80–84, 85–89, or ≥90 years), sex (female or male), educational level (less than primary or primary or more), length of stay in the nursing home (<3 or ≥3 years), dementia (yes or no), number of chronic conditions other than dementia (0−2 or ≥3), and functional dependency (no/mild, moderate, or severe/total).

**Table 2.**
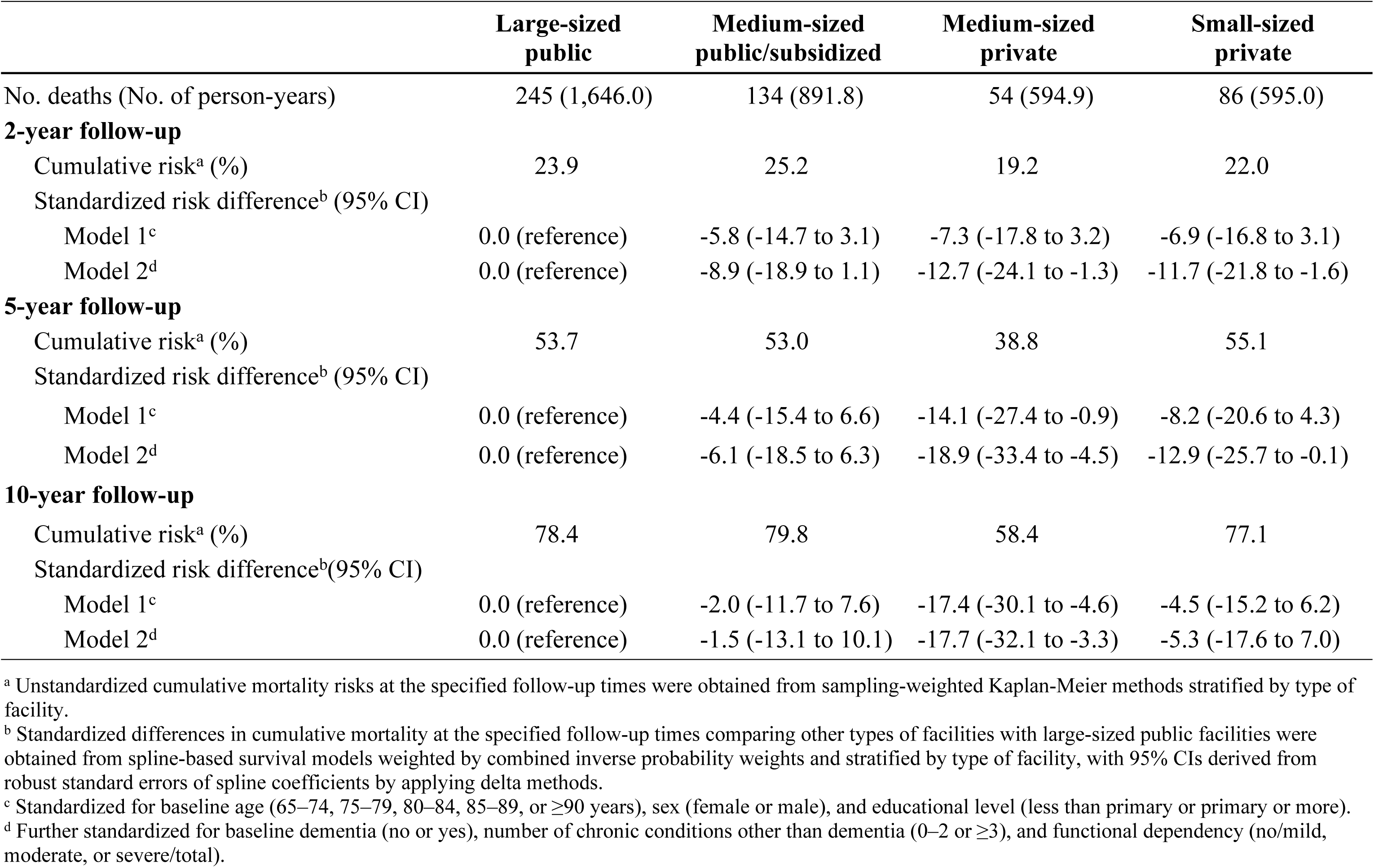
Standardized differences in cumulative all-cause mortality at 2, 5, and 10 years of follow-up by type of facility among institutionalized residents in Madrid, Spain 1998–1999 to 2013.

**Table 3.**
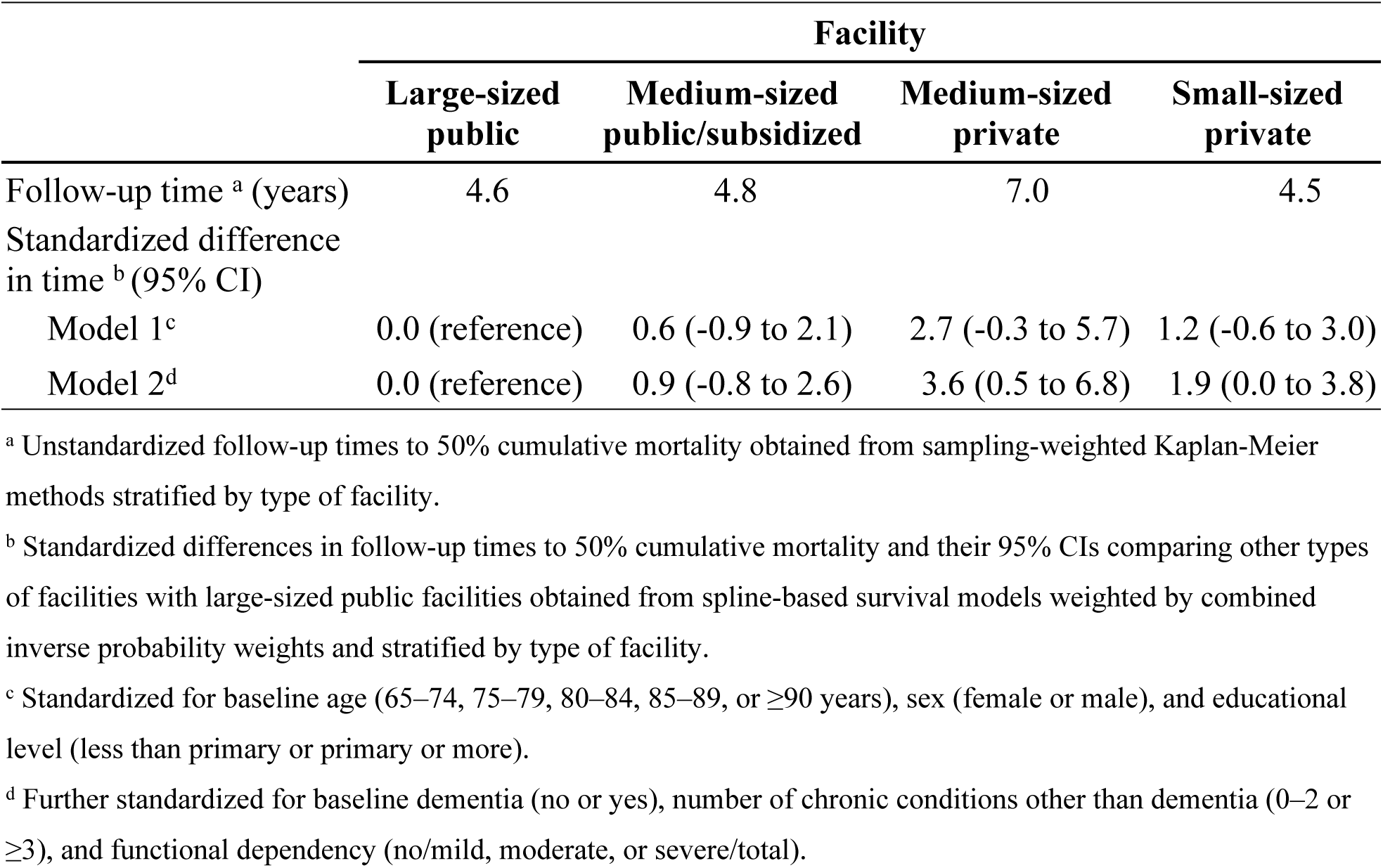
Standardized differences in median survival time by type of facility among institutionalized residents in Madrid, Spain, 1998–1999 to 2013.

Fig 2 shows results on subgroup analyses. Mortality differences among types of facilities were larger in residents with dementia, with standardized 5-year risk differences for residents with dementia of −11.9% in medium-sized public/subsidized, −42.1% in medium-sized private, and −37.3% in small-sized private facilities compared with those in large-sized public institutions.

**Fig 2.**
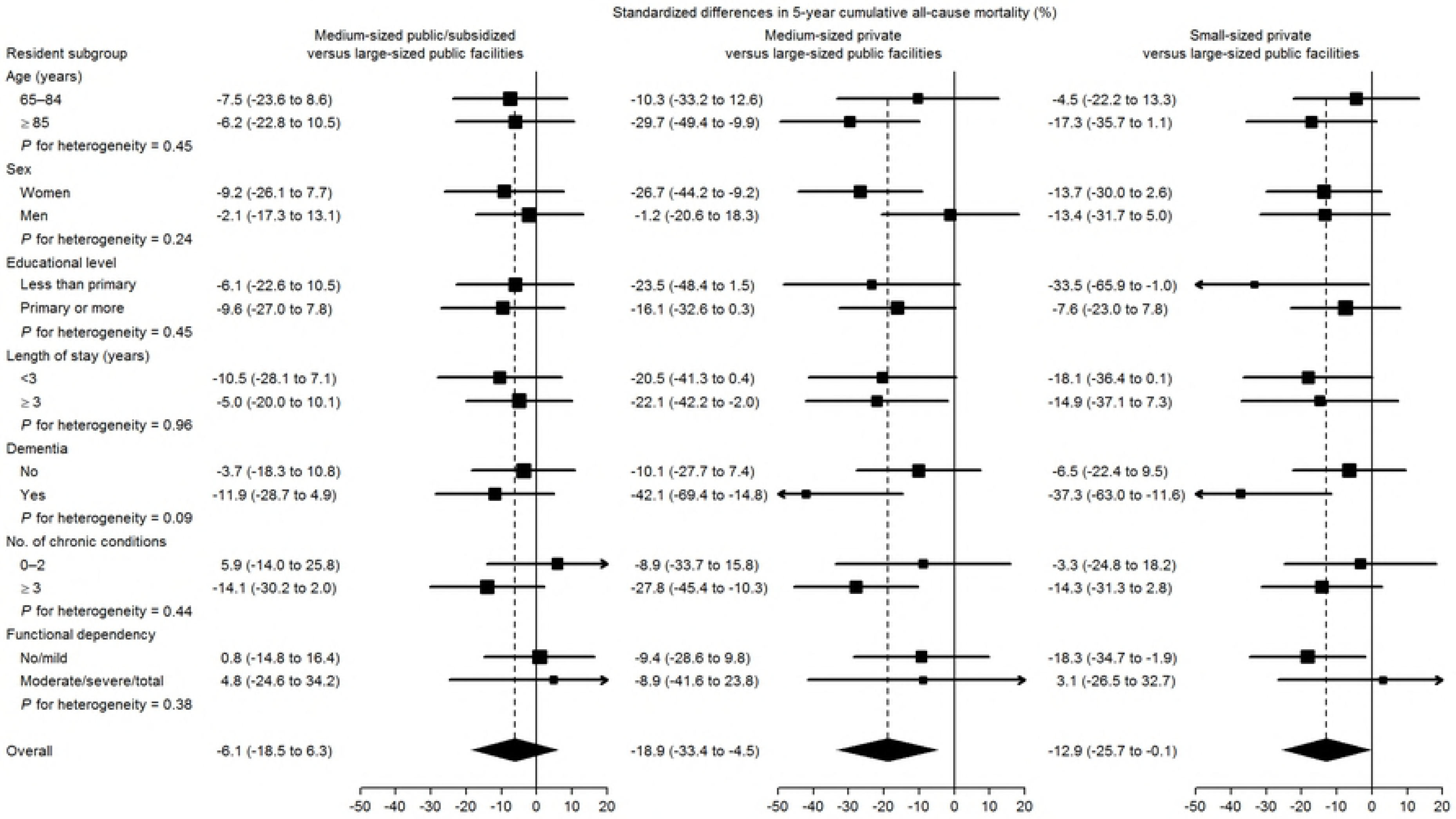
Standardized differences in 5-year cumulative all-cause mortality, (reference: large-sized public facilities) in pre-specified subgroups. Subgroup-specific risk differences (squares with area inversely proportional to the variance) and their 95% CIs (horizontal lines) were obtained from spline-based survival models weighted by combined inverse probability weights and stratified by type of facility and resident subgroup. Subgroup-specific weights were used to standardize cumulative mortality in each type of facility and resident subgroup to the weighted distribution of baseline confounders in the entire resident subgroup, including age (65–74, 75–79, 80–84, 85–89, or ≥90 years), sex (female or male), educational level (less than primary or primary or more), length of stay in the nursing home (<3 or ≥3 years), dementia (yes or no), number of chronic conditions other than dementia (0–2 or ≥3), and functional dependency (no/mild, moderate, or severe/total).

In secondary analyses according to facility ownership and profit status, the fully-standardized 5-year mortality differences (95% CIs) comparing for-profit subsidized, not-for-profit private, and for-profit private facilities with not-for-profit public institutions (as reference) were −1.4% (−20.2% to 17.5%), −9.7% (−22.3% to 2.9%), and −15.1% (−31.1% to 0.9%), and the fully-standardized median survival differences (95% CIs) were 0.2 (−2.1 to 2.5), 1.3 (−0.4 to 3.1), and 3.0 (−1.7 to 7.7) years, respectively.

## Discussion

Our study provides novel information on mortality risk not only in relation to ownership, but also in terms of its long-term perspective. Moreover, we present absolute measures of association, which facilitates the appraisal of the public health potential impact. We found a clear association between ownership type and mortality, with private facilities showing a lower risk. In addition, the association was notable in absolute measures: in five years, for every 100 residents of large public facilities an excess of 19 deaths is expected compared to what would happen if they were in medium-sized private residences. Or equivalently, the number needed to harm is more than five (1/0.19). Looking at survival time provides another way of appraising this important effect: the median life-expectancy was 3.6 years higher in the medium-private group compared to large-public. We also found a lower mortality associated with private for-profit status.

### Literature findings (see also Table S1 at supplementary material)

Spector and Takada 1991 [4], in a sample of 2500 residents from 80 nursing homes in Rhode Island, USA, found the six-month probability of death to be 58% lower in for-profit facilities. However, we consider this follow up time as too short. Bell & Krivich [5] found, in an ecological study on Illinois (USA) nursing homes, an adjusted 3% lower mortality rate in for-profit facilities compared to government and not-for-profit facilities. Deaths were recorded only while occurring in the nursing home, thus implying a probable underestimation, with this bias affecting for-profit facilities more because of a stronger tendency in for-profits towards discharging the terminally ill to hospitals, the authors said. In another ecological study Zinn et al [6] also showed for-profit status associated with lower than expected mortality in Pennsylvanian nursing homes. Cohen & Spector [7], in a sample of 2663 residents from 658 nursing homes, studied the effect of different reimbursement types on outcomes and found lower, though virtually null, mortality in public and not-for-profit compared to for-profit. Spector, Selden & Cohen [8], in a representative sample of 2230 residents from private facilities, showed results, broken down by Medicaid coverage, on the association between not-for-profit status and mortality. Mortality appeared to be slightly higher in not-for-profits among Medicaid-covered residents, but clearly lower in private-pay residents, who represented only 28% of the sample. Bliesmer et al [9] found a lower risk of dying in newly-admitted residents in Minnesota for-profit facilities, in a study analyzing three 1-year follow-up cohorts. In a large 3-year follow-up study, Porell et al [10] found 6% lower mortality in not-for-profit Massachusetts Medicaid residents. Intrator, Castle & Mor [11] found practically no difference between for-profit and not-for-profit residents’ 6-month mortality from a sample of 2080 USA residents from 253 nursing homes. In a Canadian nursing home population McGregor et al [12], found 18% higher crude mortality rate in not-for-profit facilities, but was null after adjustment for age group, sex, level of care, previous hospitalization and facility size. Finally, in 640 publicly-funded Canadian facilities, Tanuseputro et al [13], report for-profit facilities as having higher rates of mortality in newly-admitted residents and at 1-year follow-up.

In view of these results it is not possible to conclude whether for-profit facilities behave better than not-for-profit regarding mortality. Besides, two reviews concluded that for-profit nursing homes appear to provide lower quality of care in many areas of process and outcome, including mortality [22, 23]. None of the reviewed studies focused results on the private/public distinction. Yet not-for-profit can be public or private, adding another dimension that can also play a role. In fact we found that not-for-profit/private facilities showed lower mortality as compared to not-for-profit/public. In addition, we found a notable difference in mortality risk between for-profit/private and not-for-profit/public facilities.

The higher mortality associated with public facilities should be reliably attributed to differences in facility characteristics (structure and process). There would be confounding if there were a greater tendency of people with worse prognosis to enter into public facilities as compared to private ones. However, we have adjusted for very strong determinants, such as functional dependency, dementia, and number of chronic morbidities. While adjusting for these important prognostic variables may account for most of the confounding, socioeconomic factors may still have a role, though adjustment for education might have partially controlled this potential source of confounding. Consequently, we think that most of the effect must be attributed to environmental (as opposed to personal) factors associated with ownership. Unfortunately, we have not collected data on these. Although we partially controlled for the effect of facility size, it was not possible to do so satisfactorily. This was because there are not very large-sized private facilities in our area, and all small facilities are private. For the same reason, we could not properly interpret the effect of size. Nonetheless, our results are compatible with a higher risk in large facilities. In fact, some of these were very large, with more than 600 beds, and perhaps this considerable size may have had an impact on the mortality risk. Finally, we believe that a group of subtle – and admittedly speculative – factors are worth commenting on. In many nursing homes the proportion of residents for whom prolonging life is questionable may be high. Holtzman and Luire found decreased aggressiveness of care to be associated with higher mortality rate [24]. One might ask if there could be a tendency to extend life at all costs, including at the expense of losing quality of life; or, on the other hand, if not prolonging life is taken somewhat more strictly than is desirable, and if these extreme tendencies might manifest themselves differently between private and public, for-profit and not-for-profit enterprises, and to what extent. All these issues are undoubtedly complicated but, in our view, it is time for them to be addressed seriously.

Positive aspects that we propose to be considered are the use of a representative sample with a very high response-rate and the participation of various types of resident, features which enhance our study’s external validity. The appreciable sample size of the study limits random error and the availability of relevant variables confers special importance on a kind of study where confounding control is paramount. In this sense, we used methods that manage to emulate a random assignment of study participants to each of the 4 types of residence and can more realistically explain the potential effect of ownership, while controlling for important individual-level determinants. In addition, some of the most significant potential problems of these methods have been overcome, with a set of weights centered at 1.00 and with no extreme values, thus with no sign of the positivity assumption being violated. Though the set of adjustment variables included the most important determinants of mortality in this population, some degree of residual confounding is still possible. It should, however, be borne in mind here that adjusting for relevant and well-measured covariates, as we did, could improve control of confounding, since these variables may collectively serve as proxies for unmeasured factors [25].

Some features may limit interpretation of the results. Our data do not permit the satisfactorily disentanglement of the effect of size from ownership, but we suspect that an important part of the effect associated with public facilities comes via their very large size. Though ownership and for-profit status has a clear correlation, for-profit is a variable that can have a particular role apart from ownership. As for the outcome, we believe that some deaths might not have been reported, something that would eventually generate non-differential misclassification and, in general, lead to underestimation of the associations.

## Conclusions

Our results lend reasonable support to the idea that differences between public and private facilities may have an important influence on mortality risk. It is possible that very large facilities may not be the best configuration in relation to outcomes. Further investigation is thus needed to confirm this finding and to elucidate which factors are relevant in explaining the differences. In addition to the for-profit/not-for-profit classification, we believe that whether a facility is public or private may also provide useful information in the study of health issues in this sector.

## Acknowledgments

We gratefully acknowledge the contribution of all residents and staff from the participating facilities.

Disclaimer. This article presents independent results and /or research. The views expressed are those of the author(s) and not necessarily those of the Instituto de Salud Carlos III.

